# Spatial Phylogenetics with Continuous Data: an Application to California Bryophytes

**DOI:** 10.1101/2024.12.16.628580

**Authors:** Matthew M. Kling, Ixchel S. González-Ramírez, Benjamin Carter, Israel Borokini, Brent D. Mishler

## Abstract

Spatial phylogenetics is premised on the idea that species are not discrete categorical entities but instead lie on a hierarchical evolutionary continuum that contains rich biological information valuable for quantifying spatial biodiversity patterns. Yet while spatial phylogenetic approaches use quantitative information to represent phylogenetic patterns, most have continued to rely on methods that discard valuable information about spatial patterns by converting continuous variables into binary categories. This includes representing geographic ranges using binary presence-absence data, classifying statistical significance into categories, and quantifying biogeographic gradients into discrete regions. In this paper we show how a full suite of spatial phylogenetic analyses, including analyses of alpha and beta diversity, neo- and paleo-endemism, biogeographic hypothesis testing, and spatial conservation prioritization, can be implemented with “smooth” methods that never remove information content by categorizing continuous data. Our analysis focuses on the bryophytes of California, an understudied group in a global plant biodiversity hotspot. Using a time-calibrated phylogeny and species distribution models for 548 species of mosses and liverworts, we profile the evolutionary diversity, compositional turnover, and conservation value of bryophyte communities across the state. Our results highlight important patterns in the diversity of this key plant group, while our methods can serve as a model for future studies seeking to maximize the information content of spatial phylogenetic analyses.

## Introduction

The field of spatial phylogenetics provides a set of methods that account for evolutionary relationships among species when quantifying spatial biodiversity patterns. This approach offers a number of advantages over species-based approaches. For example, it better represents the genetic and phenotypic diversity present in a given area, it highlights macroevolutionary patterns like areas of neo- and paleo-endemism, and it is robust to arbitrary and inconsistent definitions of what constitutes a species (Faith 1992; Mishler et al. 2014; Tucker et al. 2018). As the field of spatial phylogenetics has matured, it has expanded to include methods for incorporating phylogenetic information into a range of analyses including inferences about alpha phylogenetic diversity and endemism, beta diversity and regionalization, biogeographic hypothesis testing, and spatial conservation prioritization (Mishler 2023). However, while these spatial phylogenetic approaches incorporate rich continuous information about the branch lengths of the phylogeny, they often neglect to take similar advantage of continuous quantitative information about spatial patterns. In a typical analysis, there are often one or more points at which information is discarded by converting continuous quantitative data into discrete categories. Here we briefly highlight three major categories of information loss that can add imprecision and bias to results. These can occur together or separately.

The first category of information loss concerns the spatial distribution data used to calculate diversity metrics. Phylodiversity metrics are almost always based on binary presence-absence data representing occurrences of the terminal taxa and larger clades (or more rarely, on abundance data; Cadotte et al. 2010). Binary data can indicate the presence or absence of occurrence records in a given spatial area, or they can be predictions from species distribution models (SDMs) used to correct for poor coverage and biased sampling in the occurrence data by estimating species’ spatial and environmental affinities. SDMs produce continuous outputs, but these have almost always been thresholded to produce binary range maps before phylodiversity analysis (e.g., Spalink et al. 2018; Allen et al. 2019), which discards information about occurrence probability. Beyond reducing precision, this thresholding can also bias diversity estimates made by combining SDMs (Dubuis et al. 2011); for instance, if a large number of taxa had a somewhat small occurrence probability in a particular site, they could all get classified as absent and the site declared low in diversity, even if their combined probability mass were higher than the probability mass in a site with only a few high-probability taxa.

A major reason for this reliance on presence-absence data is that many phylodiversity measures were originally conceived to operate on binary data. Most implementations of measures like phylogenetic diversity (PD; Faith 1992), phylogenetic endemism (PE; Rosauer et al. 2009), relative phylogenetic diversity (RPD), and relative phylogenetic endemism (RPE; Mishler et al. 2014) have used presence-absence data. However, it is also possible to use probabilities when calculating these phylogenetic measures (Kling et al. 2018). A probabilistic approach requires two modifications to the binary approach. First, occurrence probabilities for the tips of the tree are used to calculate occurrence probabilities for every clade on the tree, representing the likelihood that at least one terminal in a clade is present in a site. Alternatively, distribution models could be fit for every clade in addition to the terminal taxa. Next, these occurrence probabilities are used as weights when calculating diversity measures, giving a more complete picture of diversity patterns than is possible with binary data.

A second category of information loss can occur during statistical significance testing. Spatial phylogenetic studies often perform statistical tests by using randomization algorithms that permute species occurrence patterns, generating a null distribution against which observed diversity metrics can be compared to assess significance. Many of these algorithms only support binary occurrence data, requiring users to convert quantitative data to binary format before significance testing, while many algorithms for quantitative data support abundance data but not continuous probabilities. However, there are several algorithms available for quantitative probability data (Peres-Neto et al. 2001; Gotelli and Ulrich 2012), which should be used when working with these data to avoid information loss.

Regardless of the input data and randomization algorithm used for significance testing, studies often discard additional information by thresholding the resulting significance values into categories of statistical “significance” and “non-significance.” This follows classical treatment of p-values in frequentist statistics, which is increasingly understood to be arbitrary and oversimplified (Wasserstein and Lazar 2016; Chen et al. 2023). By instead presenting the continuous significance values, practitioners can preserve useful information representing the degree of evidence that a particular result is nonrandom. For example, in this paper we demonstrate how CANAPE (Categorical Analysis of Neo- And Paleo-Endemism; Mishler et al. 2014), which involves categorizing significance values for PE and RPE, can be modified to remove any thresholding. We use the term “SNAPE” (Smooth Neo- And Paleo-Endemism analysis) to describe the combination of continuous occurrence probability data from SDMs, quantitative null model analysis, and a continuous visualization approach.

Thirdly, a final area in which useful continuous information is often lost is when studying regional patterns of turnover in phylogenetic community composition. A common practice is to identify phylogenetic regions by performing a cluster analysis that categorizes sites into a small number of groups (Daru et al. 2017). Clustering is a valuable tool to explore biogeographic structure, and can be robust in cases where sites genuinely fall into distinct groupings. But in commonly observed situations when community composition follows a relatively smooth gradient characterized by continual turnover rather than sharp biogeographic breaks, clustering can be misleading. The alternatives are not perfect, since the underlying data are high-dimensional (with one dimension per node in the phylogeny) and visualizing it inevitably involves information loss. But approaches like ordination that embed the data in low-dimensional continuous space and visualize it using multidimensional color palettes (Ferrier et al. 2007) can help represent the continuous nature of the biogeographic gradients in ways that clustering cannot.

In this paper we demonstrate, in a range of spatial phylogenetic analyses, how continuous data can be fully retained to minimize information loss in each of these three areas. None of the methods we use is wholly new, but here we bring them together for the first time in an integrated set of quantitative approaches for spatial phylogenetics.

Our empirical analysis focuses on the bryophytes of California. California has served as a model system to study plant spatial phylodiversity thanks to the depth of its floristic information and its importance as a global plant biodiversity hotspot (Baldwin et al. 2017; Thornhill et al. 2017; Kling et al. 2018). But the bryophytes—three diverse clades of nonvascular plants including mosses, liverworts, and hornworts—have often been overlooked in biodiversity and conservation studies in California and elsewhere. Spatial occurrence data is relatively sparse for bryophytes, likely due in part to their being small, inconspicuous, and sometimes challenging to identify. For example, the Global Biodiversity Information Facility (GBIF) includes ∼40 times more records for vascular plants (tracheophytes) than for bryophytes, in spite of the two groups co-occurring in most locations. This supports bryophytes as a priority for filling in gaps in our biodiversity knowledge, a realization that has motivated recent efforts to document the bryological diversity of this California (see, for example, Shevock 2015, 2021). It also makes bryophytes good candidates for species distribution modeling and other approaches that maximize use of limited spatial information.

In this paper we develop a new dataset comprising a time-calibrated phylogeny and occurrence data for 548 species of mosses and liverworts across California (68% of species currently recognized in the state), and employ the continuous approaches described above to develop a portrait of bryophyte biodiversity across the state. This paper has three goals: (1) To explore continuous measures of phylodiversity, including the new SNAPE method. (2) To characterize the phylodiversity of bryophytes across California, including both alpha and beta diversity patterns. (3) To apply a conservation prioritization algorithm previously used for vascular plants (Kling et al. 2018) to identify regions of priority for bryophyte conservation and to compare them to those for vascular plant conservation.

## Methods

### 1. Occurrence data

All available herbarium specimen records for California bryophytes were downloaded 9 Dec 2023 from the Consortium of Bryophyte Herbaria (https://bryophyteportal.org/). Infraspecific taxa were collapsed to binomials, and synonyms were resolved using a custom thesaurus developed from the taxonomic literature and information from the TROPICOS database (https://www.tropicos.org). A current, vouchered list of accepted species documented from California was obtained from J. Shevock (California Academy of Sciences, pers. comm. December 2023), and specimens with names excluded from the California list (after standardizing nomenclature) were removed from the dataset. Species concepts and nomenclature generally followed treatments in the Flora of North America (Flora of North America Editorial Committee, eds. 1993+) for mosses, and Stotler and Crandall-Stotler (2017) for liverworts, with exceptions in taxa that have been revised since publication of those sources.

Data cleaning included removal of duplicate specimens and specimens lacking geographic coordinates. Specimens for which geographic coordinates did not fall within the county on the specimen label (with a 10 km buffer) were also removed. The initial download comprised 146,546 records, of which 63,003 were retained after cleaning (see Fig. S1a for map of these). Given the extremely low density of hornwort records, we discarded this lineage of bryophytes for downstream analyses.

### 2. Phylogeny inference

We searched and downloaded genetic sequences for six commonly used markers (*rbcL*, *matK*, *trnK*, *trnL*, *ndhF,* and *ITS*) for all the species present in the occurrences dataset using the R package ‘rentrez’ (Winter 2017). After checking the availability of genetic data, we selected four markers for liverworts (*rbcL*, *trnK*, *trnL*, and *ITS*), and five markers for mosses (*rbcL*, *matK*, *trnK*, *trnL*, and *ITS*) that maximize the representation of taxa while minimizing the amount of missing data. All the markers were individually aligned using MAFFT v.7.490 (Katoh et al. 2002) using the ‘--adjustdirection’ setting, and gene trees were constructed using IQtree2 v.1.6.12 (Minh et al. 2020) to scan for abnormally long branches, which are often due to misalignment resulting from marker or species misidentifications. After removing those sequences, we inferred species trees—for liverworts and mosses independently—using a concatenated alignment partitioned per marker, where each partition was assigned a GTR + G + I model. All the trees were inferred using IQtree2, and branch support was assessed using 1000 iterations of rapid bootstrap (Minh et al. 2020).

The phylogenetic trees were calibrated to time using secondary calibrations from Bechteler et al. (2023). For both liverwort and moss phylogenies, we selected six calibration nodes including the root (Table S1) and used a relaxed clock model specified in the ‘chronos’ function in R package ‘apè (Paradis et al. 2004). In order to obtain a single tree to perform combined “liverwort + moss” downstream analyses, we manually connected the two chronograms at their roots using the mean estimate of divergence time between mosses and liverworts (486 Mya) reported by Bechteler et al. (2023). See Fig. S1b for the tree used.

Of the 805 species with spatial data, 68% had available genetic data enabling their inclusion in the phylogeny. These 548 focal species, including 443 moss species and 105 liverwort species, were used in all subsequent analyses.

### 3. Species distribution models (SDMs)

The occurrence dataset included 51,171 records of the 548 focal species across California (Fig. S1a). Species distributions were modeled as a function of three properties: climatic suitability, landscape intactness (Fig. S2), and proximity to occurrences. These SDM methods closely followed those used in an analysis of California vascular plant diversity by Kling et. al (2018), ensuring that our bryophyte biodiversity patterns would be comparable with those reported patterns for vascular plants. Distributions were modeled within California at a 810 m spatial scale.

Climatic suitability was modeled as a function of four climate variables: climatic water deficit, log total annual precipitation, average summer maximum temperature, and average winter minimum temperature. We used mean climate data from the California Basin Characterization Model (Flint and Flint 2014) for the 1951–1980 time period. We fit a Maxent niche model (Dudik et al. 2004) for each species, using only linear and quadratic features. Target-group bias correction (Barber et al. 2022) was used to correct for spatial sampling, using background data on herbarium specimen density for all vascular plants to represent the intensity of botanical occurrence data collection efforts. With the same climate data, fitted models were then used to make suitability predictions for each grid cell across California.

Proximity to occurrences was included in order to avoid predicting species distributions in sites that are climatically suitable but far from known species occurrences. Proximity was modeled as a function of the distance between a grid cell and the nearest occurrence record for a given species. We used a half-Gaussian distance-decay function with a standard deviation of 15 km, scaled so that proximity values decreased from 1 in grid cells with occurrences, asymptotically approaching 0 with increasing distance.

We used the same landscape intactness data as was used in Kling et. al (2018). Beginning with published data on California ecological intactness (Degagne et al. 2016), we rescaled values to the 0–1 range and resampled them to our 810 m modeling grid using bilinear interpolation. This downweights sites that are ecologically degraded due to urbanization, agriculture, or other human activities, reflecting decreased probability of occurrence for most species.

These processes produced scores for a given grid cell for each of the three sub-models (climatic suitability, geographic proximity, and landscape intactness), which we multiplied to derive a final occurrence score between 0 and 1. To preserve the full information content of the modeled distributions, instead of thresholding these continuous values to produce binary presence-absence maps, we used the continuous values throughout the biodiversity analyses described below.

Calculating the spatial phylodiversity measures described below requires data on the spatial distribution of every clade on the phylogeny, not just the terminals. As implemented in the ‘phylospatial’ R package (Kling 2024), for a given clade in a given grid cell, this is calculated as 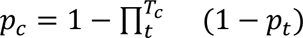, where *p_c_* is the probability of at least one terminal in clade *c* occurring in the cell, *T_c_* is the set of terminals in clade *c*, and *p_t_* is the probability of terminal taxon *t* occurring in the site.

### 4. Alpha diversity

Alpha and beta phylogenetic diversity metrics were computed using the ‘phylospatial’ (Kling 2024) and ‘canaper’ (Nitta 2021) R packages. We calculated several alpha diversity measures including terminal diversity (TD, equivalent to species richness in cases like ours where all terminals are species), terminal endemism (a.k.a. weighted endemism), phylogenetic diversity (PD), phylogenetic endemism (PE), relative phylogenetic diversity (RPD), and relative phylogenetic endemism (RPE). All metrics used the output of the SDMs as a continuous variable. For example, whereas PD with binary data is calculated as the sum of the branch lengths connecting all taxa occurring in a given grid cell to the base of the tree, PD with continuous data is calculated as the sum of the product of every clade’s branch length and its probability of occurring in that grid cell (*p_c_* in equation above). See Table S2 for equations for the derivation of all continuous diversity measures.

To assess statistical significance of the phylogenetic metrics, a community randomization algorithm is used to generate a null distribution of values for each grid cell, against which the observed values are compared in order to identify cells that have significant properties. For our main analyses, to avoid information loss from thresholding the SDMs, we used the “quantize” method (Kling 2024), which allows continuous data; we performed 1000 randomizations using five strata on a square-root probability scale. Internally, quantize utilizes a binary algorithm for different quantitative strata of a data set; we chose the “curveball” algorithm (Strona et al. 2014) for consistency with prior phylodiversity studies and because it has desirable properties like maintaining marginal row and column sums. 10000 burn-in iterations were used for each curveball randomization. We calculated significance values for all diversity and endemism measures mentioned above.

To assess patterns of neo- and paleo-endemism, we used SNAPE, a variant of CANAPE (Mishler et al. 2014) designed to maximize information content by using continuous quantities throughout the process rather than applying thresholds. In addition to using non-thresholded SDM inputs, quantitative versions of the phylodiversity metrics, and a quantitative randomization algorithm, this approach uses a smooth two-dimensional color gradient to examine the joint distribution of significance values for PE and RPE.

### 5. Beta diversity

We analyzed patterns of bryophyte community composition and phylogenetic beta diversity using the ‘phylospatial’ R package (Kling 2024). Beginning with a phylogenetic community matrix (which includes columns for every taxon, not just terminals), we normalized all rows to unit sums in order to emphasize composition rather than relative richness. We then generated a pairwise distance matrix with values representing phyloSorensen’s dissimilarity between grid cells.

To assess whether discrete regions described the dissimilarity data well, we used visual approaches such as the “elbow” and “silhouette” methods, as well as a range of quantitative algorithms in the ‘NbClust’ R package (Charrad et al. 2014) designed to identify the optimal number of clusters in a data set. None of these approaches identified a clustering scheme that fit the data well, so we instead used a continuous approach to represent smooth gradients of community turnover.

We used nonmetric multidimensional scaling (NMDS) to reduce the distance matrix to three dimensions, which we mapped to the red, green, and blue bands of color space. This approach generates a figure in which sites with similar community phylogenetic composition are depicted in more similar colors, a technique commonly used to represent biogeographic patterns (Ferrier et al. 2007). We also performed a hierarchical clustering analysis to construct a dendrogram visualizing the similarity among sites, grouping grid cells using the UPGMA (unweighted pair group with arithmetic mean) cluster agglomeration method. While neither the NMDS ordination nor the hierarchical clustering analysis preserve the full dimensionality of the community turnover data, together they paint a much richer and more nuanced portrait of biogeographic structure than does a flat cluster analysis.

### 6. Conservation prioritization

We performed a conservation prioritization to identify optimal areas where the creation of new protected areas would maximally increase protection of bryophyte biodiversity across California. In addition to the bryophyte distribution and phylogenetic data, we also used data on the locations of existing protected areas across California, including various categories of public land and conservation easements (Kling et al. 2018). Rather than binary data that treats all forms of protected land equally regardless of land management practices, these data are continuous quantitative values reflecting the relative degree of biodiversity protection based on expert knowledge of the ownership and management priorities (Fig. S2). The prioritization takes these quantities into account when assessing taxon protection levels across their ranges and when assessing potential gains from fully protecting partially-protected sites.

We used a stepwise algorithm described in Kling et al. (2018) and implemented in the ‘phylospatial’ R package (Kling 2024), which maximizes the conservation benefit of a proposed reserve network by accounting for diversity, evolutionary uniqueness, geographic rarity, conservation status, and complementarity. The algorithm is an iterative process beginning with the existing set of protected land across the study area. At each iteration, the algorithm evaluates all sites across the study area, assigning high “marginal value” to sites that are currently poorly protected and are rich in taxa that have long evolutionary branch lengths, small geographic ranges, and occur largely outside existing protected areas and other high-priority sites. Based on these marginal values, a site is selected (using one of the two criteria described below) to add to the protected land network, it is marked as fully protected, and the next iteration of the algorithm begins. Sites marked as protected earlier in the process are considered higher priority.

We performed two variants of the optimization analysis. The first identified an “optimal” conservation solution by selecting the site with the highest marginal value at each iteration of the algorithm. The second used a “probabilistic” approach in which the probability of a site being selected is proportional to its marginal value. For the probabilistic analysis, we performed 1000 randomized runs of the algorithm, and calculated the fraction of these solutions in which each grid cell was in the top 50 highest priorities. The probabilistic approach relaxes the unrealistic assumption that protected land will actually be added in the optimal order, allowing redundant high-value sites to each be recognized as valuable on different randomized runs of the algorithm.

### 7. Contrasting continuous vs. discrete phylodiversity methods

In addition to the empirical analyses, we also explored the effects of using continuous versus discrete methods. For this comparison we focused on the neo- and paleo-endemism analysis, comparing the continuous SNAPE results described above to a traditional CANAPE analysis that involved discretization at both the SDM thresholding and p-value thresholding stages.

We converted SDM predictions to binary presence/absence maps using a threshold of 25% of the maximum predicted probability for a given species. Note that while SDM studies often use species-specific thresholds chosen to optimize predictive performance, we used an arbitrary threshold because the proximity portion of our SDM methods is incompatible with threshold optimization, and because the purpose of thresholding in this case was methodological comparison rather than predictive performance. We then used these binary data in a CANAPE analysis, using the curveball randomization algorithm and thresholded significance values. In this method, p-values are thresholded to categorize cells as significant centers of endemism (in a one-tailed test of PE), and cells passing this first test are classified as significant concentrations of paleo-endemism, neo-endemism, or mixed-endemism (in a two-tailed test of RPE). In addition to the “full CANAPE” and “full SNAPE” approaches, we also assessed endemism using two hybrid approaches, one using binary SDM data but smooth visualization of p-values, and another using continuous SDM data but thresholded categorization of p-values.

All spatial analyses were done in R (R Core Team 2023). Code, spatial data, DNA alignment + tree file, and intermediate results are available at https://github.com/matthewkling/ca_bryo_sphy.

## Results

### 1. Diversity and endemism patterns

The spatial pattern of PD closely followed that of terminal diversity (TD, i.e. species richness), with higher values in the northwestern part of the state and extending south along the Pacific coast and the Sierra Nevada Mountains. Randomization results indicated that both PD and RPD are significantly high along the northern coast of the state, and significantly low in arid regions of southern and southeastern California (Fig. 1). While PE and PE significance were also highest in the northwest and lowest in the southeast parts of the state, PE exhibited a patchier and more variable spatial pattern than PD, including numerous pockets of high PE in locations with relatively low PD. (Fig. 1).

**Figure 1.**
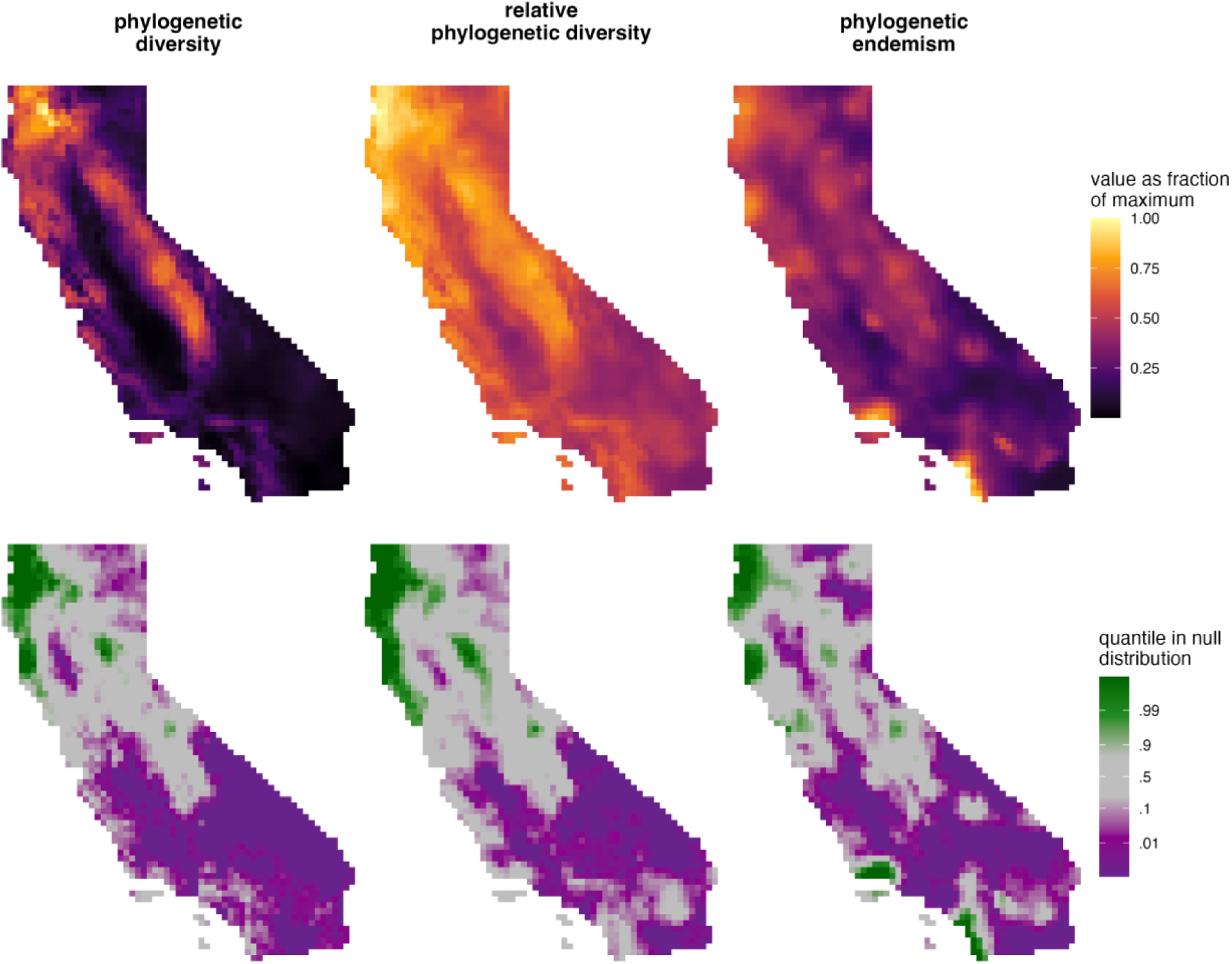
Diversity patterns of bryophytes (liverworts and mosses) in California. The first row shows PD, RPD, and PE, each scaled as a proportion of the maximum value observed across the study area. The second row shows significance values for each of the metrics based on a randomization with a null model for continuous data.

Diversity patterns for the mosses alone (Fig. S3) were very similar to these combined results. The liverworts alone (Fig. S4) showed roughly similar diversity patterns to the mosses, but had higher diversity values in the Channel Islands, more restricted areas with significantly high or low PD and RPD, and different areas with significantly high PE.

### 2. Areas of endemism significance with SNAPE

We found six zones of significant endemism when looking at the complete bryophyte dataset (liverworts + moss, Fig. 2c). The four zones located along the coast were all predominantly characterized by paleo-endemism (including the Cascades coast (*ii*), San Francisco Bay area (*iii*), Santa Barbara region (*iv*), and San Diego area (*v*)), while the two inland zones had regions characterized by mixed- and neo-endemism (including a large portion of the Sierra Nevada (*i*) and a smaller zone in the Mojave Desert near the Nopah Range Wilderness Area (*vi*)).

**Figure 2.**
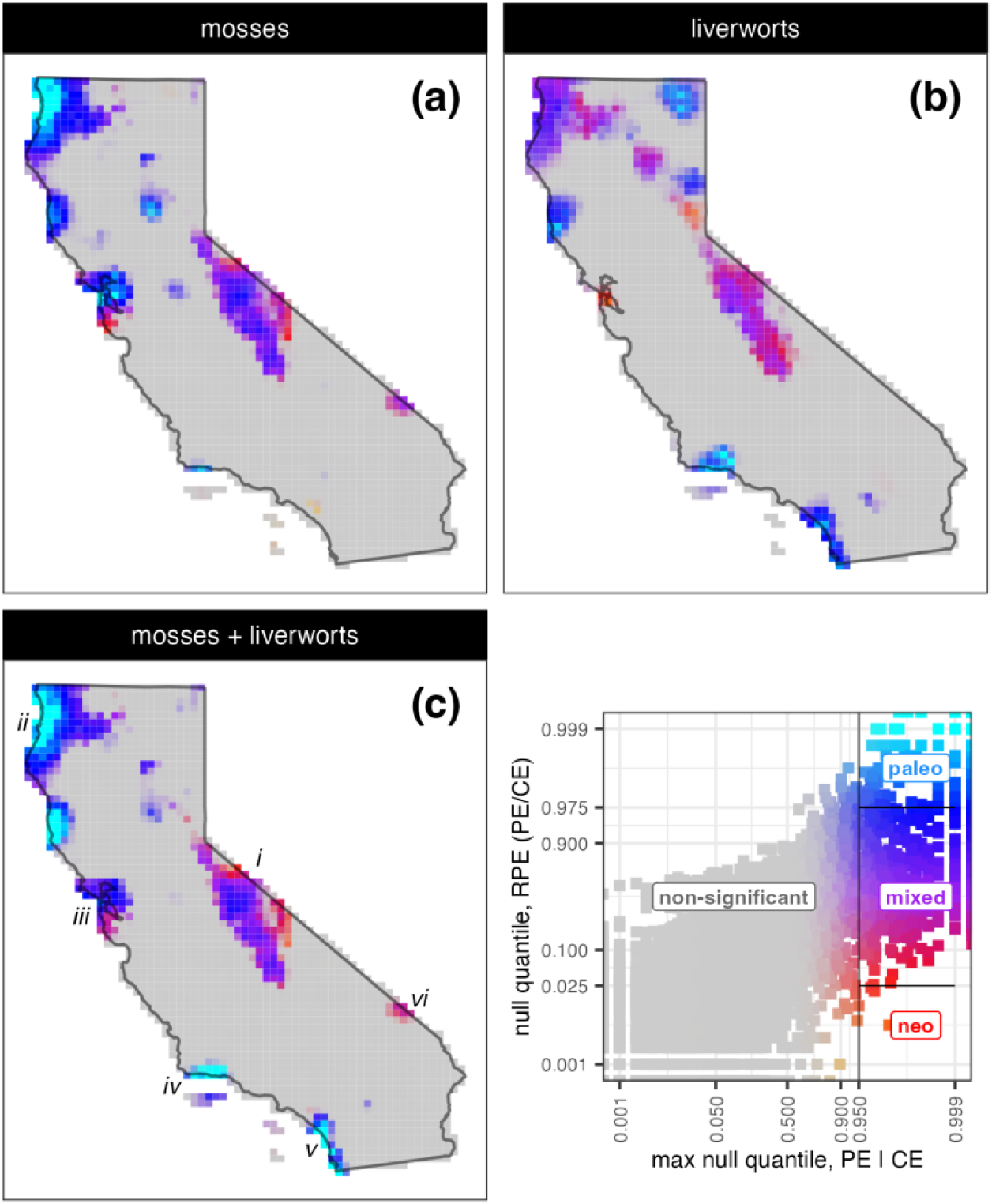
Neo- and paleo-endemism results using SNAPE for **(a)** mosses alone, **(b)** liverworts alone, and **(c)** all bryophytes. The numerals in panel **c** denote endemism zones discussed in the text. These analyses use ‘quantizè to treat SDMs as continuous variables in the spatial randomizations, and are colored following the smoothed p-value approach described in the text. Colors indicate endemism profiles defined by the combination of PE and RPE significance, as shown on the inset scatter plot. On the scatter plot, points represent grid cells on the map, black reference lines delineate the category boundaries used in CANAPE, and axis scales are logit-transformed to highlight significance magnitudes.

Endemism patterns for mosses alone (Fig. 2a) and liverworts alone (Fig. 2b) were generally similar to the endemism patterns of the combined data set, though the details differ in some important areas. Regions dominated by paleo-endemism along the southern coast (*iv* and *v*) were significant for the full dataset and for liverworts, but were reduced or missing for the mosses. The neo-endemism zone in the eastern Mojave Desert (*iv*) was significant for the full data set and for mosses, but not for liverworts alone. There was also a region dominated by paleo-endemism in the Modoc Plateau in northeastern California that was significant only for the liverworts, and a region dominated by paleo-endemism in the northern Central Valley that was significant only for mosses (Fig. 2).

### 3. Beta diversity

Our analysis identified strong patterns of bryophyte compositional differences across California. These patterns were generally consistent with major vegetation types across the state. The ordination axes separate montane regions with high TD and PD (i.e., the Sierra Nevada, Cascades, and Klamath Range) from lower elevation, drier, and more mediterranean regions that tend to have lower TD and PD (Fig. 3). Similarly, the deepest split in the hierarchical cluster analysis separates the southern deserts and San Joaquin Valley from the rest of the state (Fig. 3). Compositional turnover aligns along major climatic axes across the state, including both the strong precipitation gradient from north to south as well as the gradient from moderate coastal climates to relatively more seasonal and extreme climates inland and at higher elevations.

**Figure 3.**
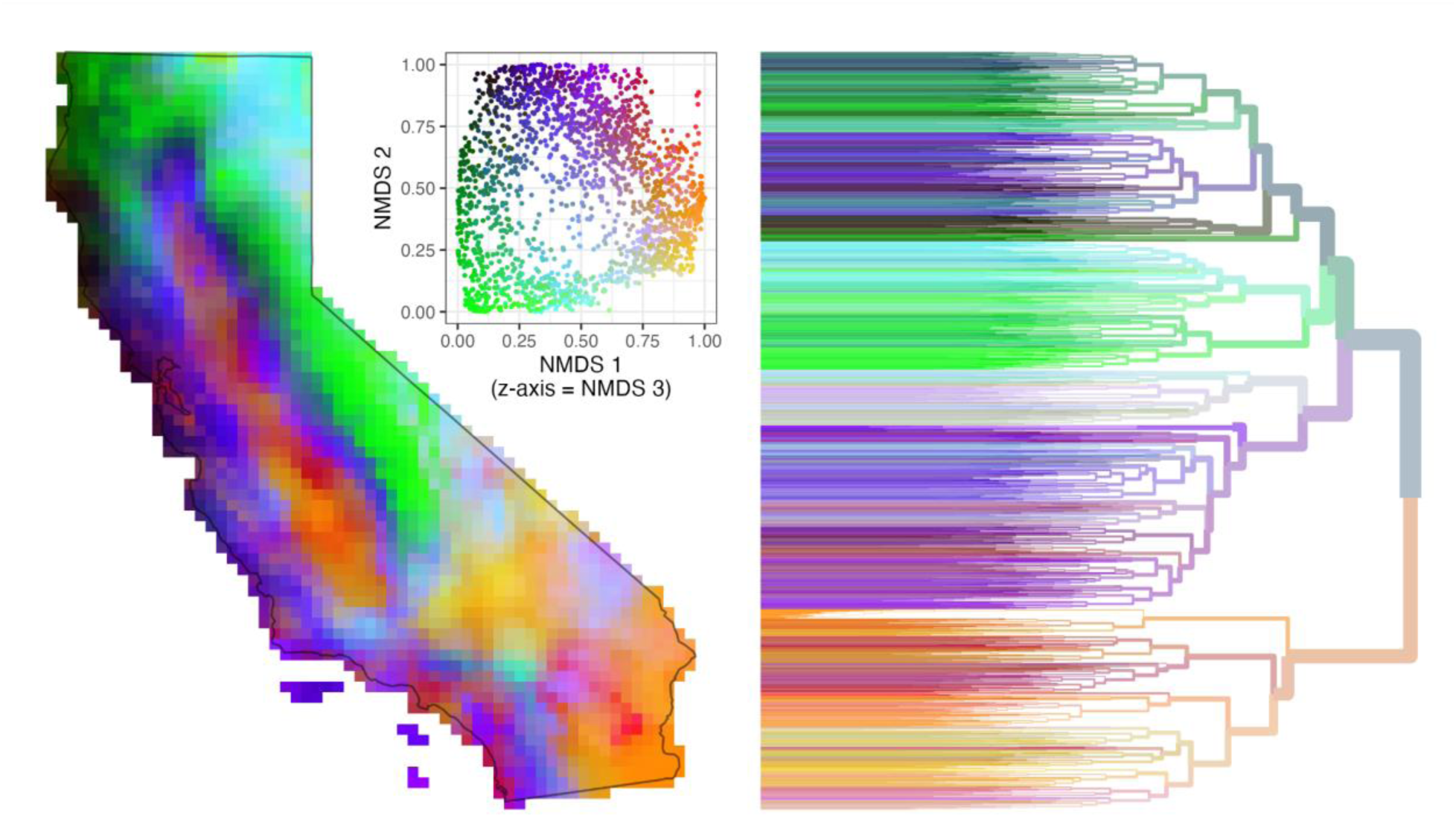
Community ordination showing compositional gradients of bryophyte communities across California. Sites depicted in similar colors have phylogenetically similar bryophyte communities. Colors are assigned based on a three-dimensional ordination. Two axes of this ordination are depicted in the inset scatter plot. The dendrogram shows a hierarchical clustering of the grid cells by phylogenetic similarity (not a phylogeny of taxa); internal dendrogram branches are shown in the average color of the cells in the cluster.

### 4. Conservation prioritization

According to the prioritization analysis (Fig. 4), the areas of highest added conservation value are primarily in the northwestern California: the land adjacent to Humboldt Bay, the coastal forests of Mendocino, parts of the San Francisco Bay area, and the area of convergence between the Sierra Nevada and the Klamath Mountains (e.g., the land adjacent to Lassen and Shasta-Trinity National forests). Other regions of relatively high interest for conservation are in the western Sierra Nevada foothills, and the Santa Barbara and San Diego coastal areas that coincide with regions of endemism in the SNAPE analyses. Note that low priority areas include regions that are relatively low in biodiversity (e.g., the southern San Joaquin Valley), as well as areas that are high in biodiversity but already well protected (e.g., the High Sierra).

**Figure 4.**
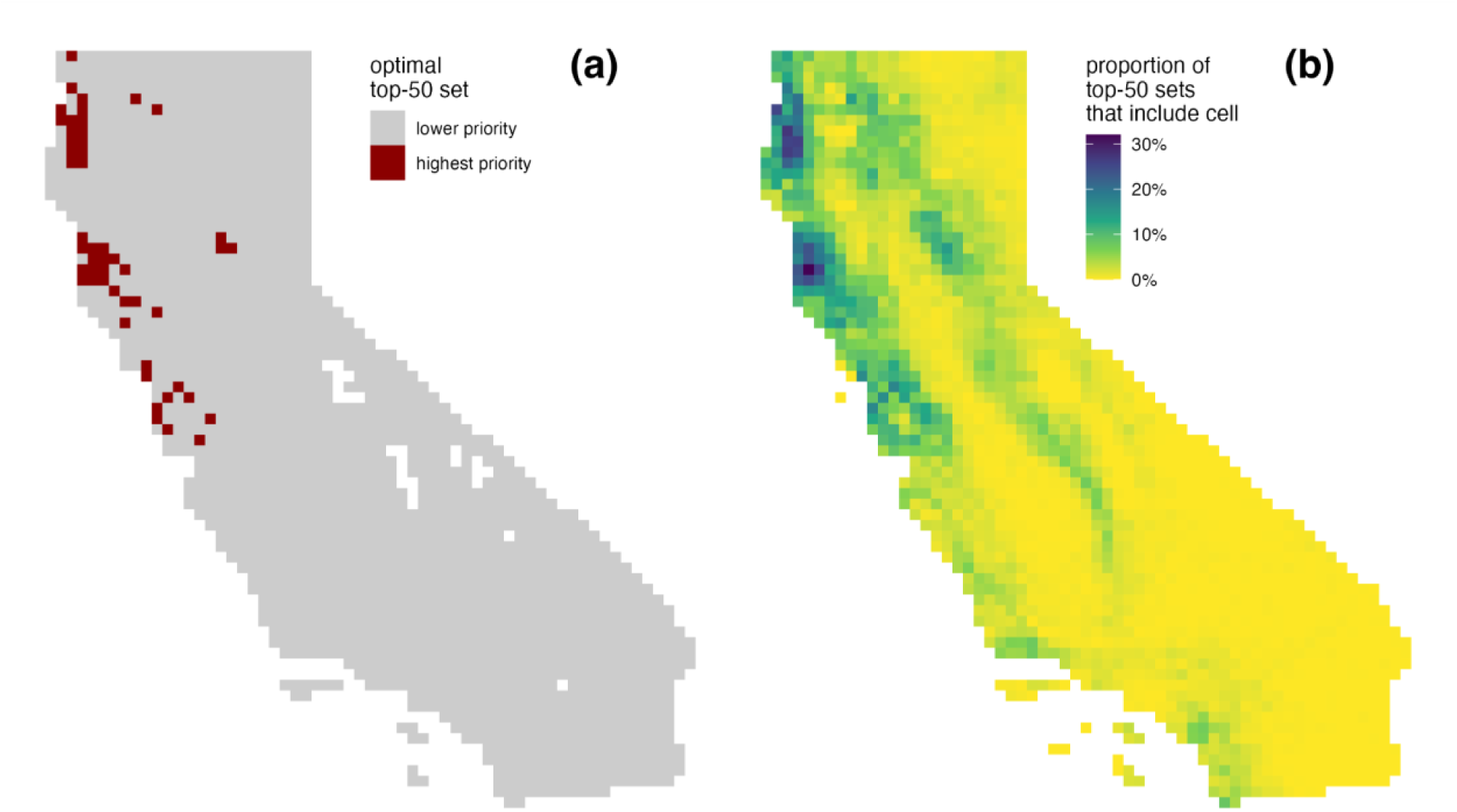
Priority locations for the creation of new reserves to increase protection of bryophyte biodiversity in California. **(a)** The top 50 highest-priority cells using optimal site selection. **(b)** The probability that a cell is in the top 50 highest priorities using probabilistic site selection. The prioritization algorithm is based on Kling et al. (2018) and accounts for diversity, evolutionary uniqueness, geographic rarity, conservation status, and complementarity of the priority areas with each other and existing protected areas. See Fig. S2 for a map of current protected areas.

### 5. Continuous versus discrete methods

We explored the effects of using continuous versus binary (thresholded) SDM distribution data, and of using smooth versus categorical interpretation of endemism significance values. Thresholding the SDMs (Fig. 5, column 1) resulted in a lack of sufficient information to categorize pixels in places where all taxa had relatively low occurrence probabilities, such as in heavily agricultural or urban areas the Central Valley or the Los Angeles Basin (white cells in Fig. 5a, 5c). In contrast, the quantitative occurrence data approach was able to analyze all cells by taking advantage of the continuous SDM output (Fig. 5, column 2). It is also notable that the isolated single-cell areas of significant endemism identified along the margins of these white no-data areas in the thresholded approach (Fig. 5a) do not appear when using the continuous approach (Fig. 5b). This suggests that the reduced information content of the thresholded SDM data is leading to these cells being randomly misidentified as areas of endemism, likely due to increased classification noise arising from higher variance in observed diversity values.

**Figure 5.**
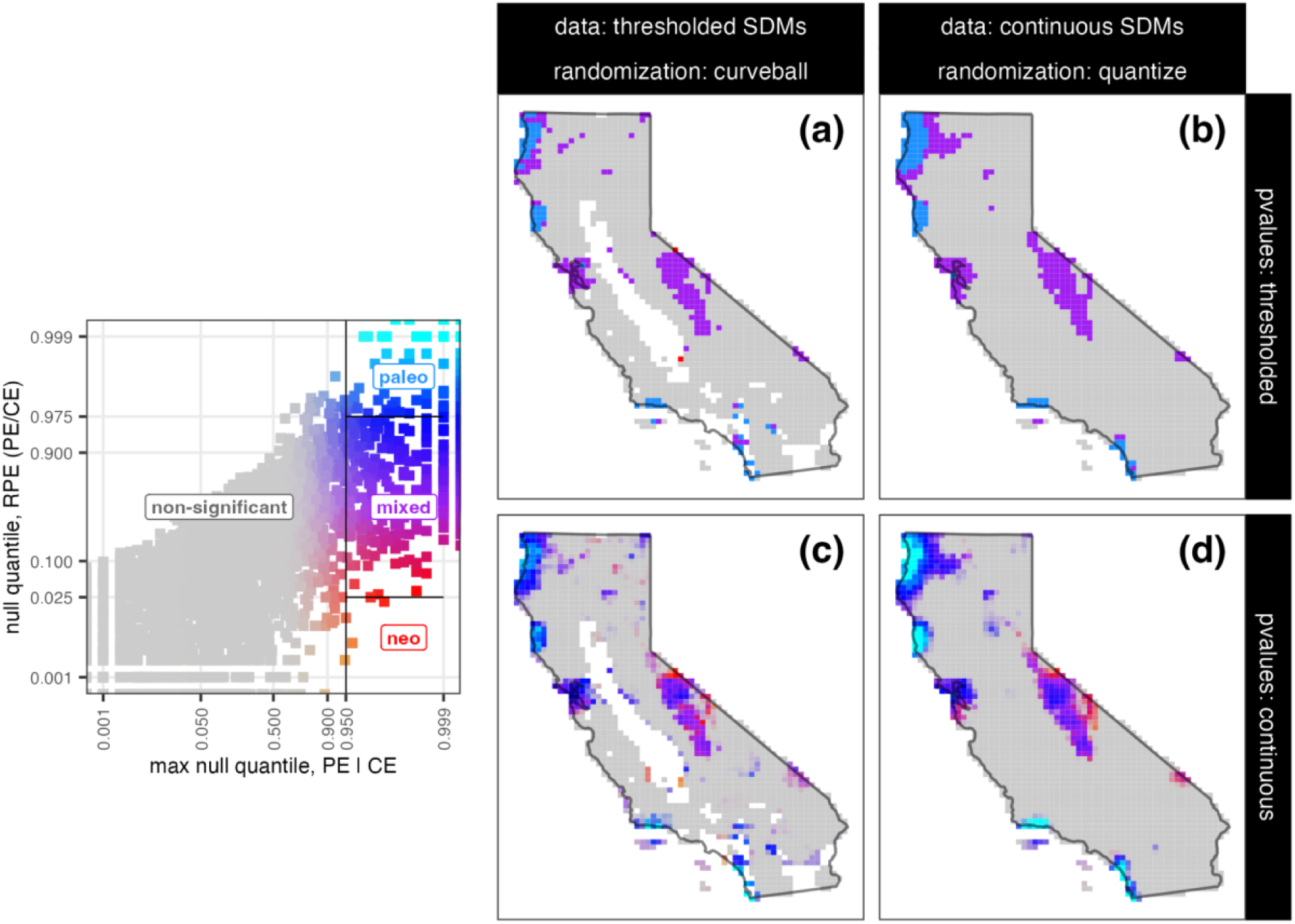
Empirical comparison of methodological approaches for identifying significant areas of bryophyte neo- and paleo-endemism. The left (**a, c**) and right (**b, d**) panels differ in whether binary versus continuous distribution data (and corresponding randomization algorithms) were used in the analysis. The upper (**a, b**) and lower (**c, d**) panels differ in whether statistical significance is communicated by classifying p-values into discrete categories versus visualizing them as a smooth surface. Panel **a** is a fully discrete CANAPE procedure, while panel **d** is a fully continuous SNAPE procedure; **b** and **c** are hybrid approaches. The legend at left shows how the mapped colors relate to p-values for PE and RPE; continuous p-values are visualized in panels **c-d** using the color gradient, while thresholded p-values are show in in panels **a-b** using only four colors corresponding to the categories delineated by the black threshold lines on the scatter plot.

Additionally, the continuous SDM approach inferred larger, more spatially continuous clusters of endemism relative to the threshold approach. For example, notice the enlarged areas of paleo- and mixed-endemism in the Cascade and Sierra Nevada regions (e.g., Fig. 5a vs. Fig. 5b and 5c vs. 5d). Most notably, the Trinity Alps region was identified as an endemism hotspot using the continuous SDMs but not the thresholded SDMs, perhaps because thresholding removed information about large numbers of taxa with a somewhat low probability of occurring there.

Use of continuous p-values highlighted structure within the major areas of endemism, and highlighted areas of “marginally significant” endemism that were not observed with the p-value threshold approach. For example, while the Sierra Nevada was inferred as a major area of endemism using any methodological approach, only the smooth approach highlighted the patterns of paleo-endemism at higher altitudes and neo-endemism in the drier eastern regions (e.g., Fig. 5b vs. 5d). Similarly, the smooth p-value approach highlighted closely co-occurring neo- and paleo-endemism regions in the Bay Area, which were categorized as mixed-endemism when the p-value threshold approach was taken (e.g., Fig. 5b vs. 5d).

## Discussion

### California bryophyte phylodiversity patterns

California has long been an important test case for understanding patterns of plant diversity and the mechanisms that underlie them (Raven and Axelrod 1978, Stebbins and Major, 1965). In recent years, this trajectory has been maintained with fine-scale analyses of taxon richness (Baldwin 2014, Baldwin et al. 2017), phylogenetic diversity, endemism, and turnover (Thornhill et al. 2017), and identification of conservation hotspots informed by spatial phylogenetics (Kling et al. 2018). In extending this effort to the state’s bryophytes, this study presents the first biogeographic analysis of California’s bryophyte flora and highlights both similarities and differences between the region’s vascular and bryophyte floras.

Note that the primary spatial phylogenetic analysis of California vascular plants by Thornhill et al. (2017) used presence/absence data from herbarium specimens of any age, thus not including any consideration of present-day intactness, and the terminal taxa were not all at the species level; in contrast, the analysis of bryophytes presented here used continuous occurrence probabilities taken from SDMs that included intactness, and the terminal taxa were all at the species level. Thus the two studies did not use entirely comparable data, yet we feel that the broad patterns of their results can be compared with caution.

Bryophytes share many traits that distinguish them from vascular plants (Mishler and Oliver 2009), and many of these traits may affect their distribution. For example, bryophytes relate to the water environment via poikilohydry and desiccation tolerance (Proctor et al 2007), as compared to vascular plants that are endohydric, which could result in different patterns of diversity in relation to moisture gradients. Other bryophyte traits, like their reproduction via spores, possibly give them high dispersal distances in comparison to seed plants. Thus one would expect differences in the spatial diversity patterns of bryophytes in comparison to vascular plants.

We found that raw TD and PD were highest in the northwestern corner of the state and were high throughout the Sierra Nevada, and extending down the coast to the Santa Cruz Mountains. Both TD and PD dropped off markedly outside these regions. This contrasts with patterns of raw TD and PD in the vascular flora (Thornhill et al. 2017; Kling et al. 2018), in which mountainous areas throughout the state had consistently high TD and PD, with marked drop-offs only in the San Joaquin Valley and in the deserts of eastern California. This difference suggests a stronger role for total precipitation in dictating TD and PD of bryophytes relative to vascular plants (Fig. S5). The striking contrasts in TD and PD patterns between these two major groups of land plants suggest that their ecophysiological differences scale up to play an important role in regional species assembly. The strong TR and PD gradient is well aligned with a moisture gradient in bryophytes but not in vascular plants. This suggests that at least locally, nestedness (depauperate regions having a subset of the species in more diverse regions) might play a stronger role in bryophytes than turnover (replacement of species along gradients), though additional analysis would be needed to confirm this hypothesis. Nestedness was inferred as being important for bryophytes in extreme northern California (Norris 1997), and was demonstrated in a comparison of bryophyte and vascular plant biogeography across the Channel Islands (Carter and Guilliams 2018). However, the broader-scale difference between the northern/coastal region and southern drylands region is likely driven more strongly by turnover (Fig. 3).

Patterns of significantly high and low PD and RPD can provide important insights into the ecological and evolutionary drivers of species assembly. Significantly low PD indicates phylogenetic clustering, while significantly high PD indicates phylogenetic overdispersion, usually interpreted in an ecological context as either habitat filtering or competitive exclusion, respectively (Mishler 2023). Significantly low RPD indicates a concentration of short branches, while significantly high RPD indicates a concentration of long branches, usually interpreted in an evolutionary context as either a center of divergence or a refugium, respectively (Mishler 2023).

The geologically older, relatively stable mesic environments of the northwestern part of the state and the northern Sierra Nevada (Raven and Axelrod 1978, Baldwin 2014) have significantly high PD (indicating that competitive exclusion could be acting) and high RPD (indicating that the region may be acting as a refugium for longer-branch lineages). On the other hand, the comparatively younger arid regions in the southern half of the state have significantly low PD (indicating habitat filtering) and low RPD (indicating relatively recent diversification of lineages adapted to arid regions) (Raven and Axelrod 1978, Baldwin 2014). The northwestern corner of California, and also higher elevations in the Sierra Nevada, harbor a relatively large number of species from early divergent moss lineages (e.g., *Sphagnum*, Polytrichaceae, Tetraphidaceae) that play a role in driving high RPD. In contrast, southern California, and especially the desert, has substantially fewer species, from a much narrower suite of lineages (e.g., Pottiaceae, Orthotrichaceae, Grimmiaceae) that are well adapted to arid conditions, many of which likely diversified relatively recently in the drylands of southwestern North America and drive the low RPD in that region.

Interestingly, the significance patterns in California bryophytes for PD and RPD were quite similar to each other (Fig. 1). This differs markedly from Thornhill et al.’s (2017) analysis of the California vascular flora, and from the general trend across other spatial phylogenetic studies that tend to reveal greater differences between PD and RPD significance patterns (e.g., Mishler et al. 2014, 2020, Thornhill et al. 2016, Scherson et al. 2017, Earl et al. 2021; Nitta et al. 2022). Considerable similarity between significance patterns of PD and RPD was also seen in a spatial phylogenetic analysis of the North American moss flora (Carter et al. 2022). That the significance patterns for PD and RPD were quite similar to each other for bryophytes in both the continental study and this study on California, but not for vascular plants at either scale, suggests that some particular aspect of bryophyte biology might be involved.

As discussed above, in mosses long-branch taxa tend to co-occur in high richness areas while short-branch taxa tend to co-occur in low richness areas (e.g., lineages adapted to the desert). When the same taxa are found together, PD and RPD vary in proportion to each other, and thus would show the same significance patterns. Thus, an unusually strong sorting of taxa along a single environmental gradient appears to explain this unusual similarity of significance patterns in PD and RPD in the mosses. In turn the pattern in the mosses drives the pattern seen for all bryophytes.

### Potential drivers of bryophyte endemism

The SNAPE analysis highlighted several regions as significantly high in phylogenetic endemism. Note that these regions are identified following the randomization, which accounts for species richness, and thus these areas of PE significance are not necessarily high in richness or PD. Below we describe each of these regions, highlighting any differences observed when considering only mosses, only liverworts, or the complete dataset including mosses + liverworts (referred to as “all bryophytes”).

*(i) The Sierra Nevada.* This was the largest area of endemism identified for bryophytes in California. Within this area, we observed variation in the predominance of different types of endemism. First, we found concentrations of neo-endemism for liverworts, mosses, and all bryophytes (Fig. 2) along the west slope of the mountains. This is likely related to the relatively recent onset of the mediterranean climate in California over the past few million years. Although not many Californian bryophyte lineages have been systematically studied phylogenetically, those that have, for example in Orthotrichaceae (Medina et al. 2012, 2013, Lara et al. 2020), and Brachytheciaceae (Huttunen et al. 2008, Carter 2012), indicate recent diversification in California. Second, there was a region of significant paleo-endemism at high elevation in the Sierra that is driven by mosses, since we do not observe high paleo-endemism for liverworts. Similar to the northwestern part of the state, the high elevation Sierra Nevada has relatively high diversity and abundance of early divergent moss lineages (e.g., *Sphagnum*, Polytrichaceae) which likely contribute strongly to this pattern.
*(ii) The Cascades.* This was the second largest area of endemism. It is located in the northwesternmost part of California, and the main vegetation consists of evergreen forests. It is a relatively wet region in comparison to the rest of the state. The SNAPE analysis categorized this area as dominated by paleo-endemism for mosses (Fig. 2b) and all bryophytes (Fig. 2c), while for liverworts it is categorized mainly as mixed-endemism. The higher contribution of neo-endemism for liverworts is likely driven by the high diversity of epiphytic leafy liverworts, a recently diversified clade (Feldberg et al. 2014), occurring in ecosystems with higher humidity. In mosses, this area has relatively high concentrations of both early divergent lineages and also long-branch terminal taxa that either without close relatives or are lineages disjunct between northwestern California and east Asia (reviewed by Carter et al. 2016).
*(iii) San Francisco Bay Area.* The native vegetation of this region includes coastal dunes, chaparral and at higher altitudes evergreen forests. It is strongly influenced by the California Current, which keeps the weather temperate all year long. While this region was not characterized as a large center of endemism for vascular plants (Thornhill et al. 2017), it was categorized as a mix dominated by paleo-endemism in the north and neo-endemism in the south for mosses and all bryophytes (Fig. 2a and 2b). Interestingly, for liverworts (Fig. 2a) only a small area in the southern part of the San Francisco Bay was found to be a center of neo-endemism.
*(iv & v) Santa Barbara and San Diego coasts.* These coastal regions of southern California are characterized by coastal sage scrub and chaparral vegetation. They were identified as areas dominated by paleo-endemism for liverworts and all bryophytes but not for mosses. This pattern is driven by the occurrence of multiple species of thalloid liverworts with very long evolutionary branches and small geographic ranges, like the highly endangered *Geothallus tuberosus*, and several species of *Riccia and Sphaerocarpos*.
*(vi) Eastern Mojave Desert*. For mosses and all bryophytes, a small area in the California desert was categorized as a significant concentration of neo-endemism. This general area was also seen as an area of high endemism in the vascular plants (Thornhill, et al. 2017), but this result for bryophytes needs to be taken with a grain of salt—it might be at least in part an artifact of low and heterogeneous collection efforts across the desert mountains.

These regions of significantly high bryophyte endemism loosely align with regions of endemism that have been defined at the scale of the North American continent. In an analysis of the North American moss flora (Carter et al. 2016) that employed only species-based metrics rather than the PE metric used here, five areas of endemism were identified in California: a California Floristic Province area, a desert southwest area which included the Mojave and Sonoran deserts in California, an interior area that included the Great Basin portions of eastern and northeastern California, a northwestern area which extended along the coast from the Aleutian Islands to near the California/Oregon border, and a Pacific Northwest area that extended from southern British Columbia in the north through the Sierra Nevada and Santa Cruz Mountains in the south. A spatial phylogenetic analysis for North American mosses (Carter et al. 2022) recovered very similar areas of endemism. Results from that study differed primarily in the deserts, with the Sonoran desert clustering distinctly away from the rest of California and with the Mojave and Great Basin desert portions of California clustering with an interior Rocky Mountain region rather than with other areas of California. Areas of significantly high endemism within California in that study were mostly areas of mixed endemism between San Francisco and the Oregon border.

### Implications for California bryophyte conservation

This is the first study we are aware of to conduct a formal spatial conservation prioritization for the bryophytes of California. At a high level, the conservation prioritization patterns we identified for bryophytes are quite similar to patterns previously identified for vascular plants (Kling et al. 2018). This is unsurprising, since the prioritization algorithm avoids existing protected areas as well as highly impacted urban and agricultural areas, regardless of the taxonomic group being analyzed. Large swaths of California fall in both these categories, leaving a relatively small portion of the state that is both unprotected and ecologically intact.

However, when we contrast the 50 sites with the highest conservation priority between vascular plants and bryophytes, we found that all high conservation priority regions for bryophytes are in northern California, unlike for vascular plants where some were in Southern California. This might be expected given that our analysis did not find many areas of significantly high endemism in the southern part of the state for bryophytes. As discussed above, these differences in endemism patterns, and thus conservation prioritization, are linked to real biological differences between these groups, such as poikilohydry in bryophytes.

It is important to note that our inferences about conservation priorities and basic biogeographic patterns were made from available herbarium records, which are incomplete and biased for bryophytes—in particular for the California desert. SDMs can help to partially mitigate these issues, by interpolating occurrence records and accounting for modeled sampling biases. We note that our implementation of SDMs did not account for micro-environmental variables, which likely play particularly important roles in many bryophyte distributions. For such species, the climatic component of our SDMs may fit poorly and produce relatively flat, uninformative climatic suitability surfaces, which results in range predictions that are informed mainly by the proximity and intactness portions of our models. Future studies could help to refine estimates of bryophyte distributions by incorporating additional predictors representing factors like finer-scale environmental variation and biotic interactions. In the longer run, another important solution is increasing the density of occurrence data by doing additional floristic work, particularly in the deserts, which have received relatively little attention.

### Avoiding information loss in spatial phylogenetic studies

These analyses illustrate that it is both straightforward and valuable to avoid information loss from discretizing continuous data in phylodiversity analyses. There are readily accessible methods for making full use of continuous data, including when representing geographic ranges, interpreting statistical significance, and visualizing community turnover. These methods are generalizable and can be easily applied to studies focused on other locations and taxa.

While the use of SDMs in spatial phylogenetics has become a common practice, the typical approach has been to transform the continuous probability output of SDMs into presence/absence data. This is in contrast to species richness studies, which have begun to make more widespread use of continuous occurrence probabilities from stacked SDMs (e.g., Dubuis et al. 2011). When SDMs are thresholded, information on the likelihood that a taxon occurs in an area—reflected in the probability value—is lost. Using “smooth” phylodiversity metrics, which weight a clade’s contribution to the diversity based on its continuous probability of occurrence, we can maximize the use of information obtained from SDMs.

When we compared endemism patterns based on continuous SDMs versus thresholded SDMs (Fig. 5), we found signs that thresholding introduced both bias and variance in the results, and also that it limited the geographic coverage of the analysis. One manifestation of bias was that some regions of the state contained no bryophyte presences after thresholding the SDMs. These were areas where all species had relatively low occurrence probabilities, including heavily agricultural or urban landscapes. SDM thresholding presumably resulted in underestimates of true diversity in these areas (assuming that at least some bryophytes almost certainly occur there, even if the probability is low for individual species). Thresholding also limited the geographic coverage of our results, making it impossible to assess characteristics of the phylogenetic community structure for these communities. In contrast, the continuous SDM analysis was able to analyze diversity and endemism for these sites. In addition to being able to utilize information on small occurrence probabilities for the terminal taxa, the continuous approach recognizes that the probability of at least one species in a large clade occurring in a site can still be quite high even if all the individual species in the clade have low occurrence probabilities. For example, under simple assumptions of independent probabilities, if a clade contained 100 species each with a 1% probability of occurring in a given site, the clade’s occurrence probability in that site would be 63%. Instead, in a binary analysis using thresholded SDMs, at least one species in a clade has to have a high occurrence probability in order for the clade to be considered present in a site—an assumption that is inconsistent with probability theory.

Our results also suggest that this form of bias may not be limited to low diversity areas. Our analyses found several relatively diverse areas, such as the Trinity Alps in inland Northern California, where analysis using continuous probabilities identified significant endemism areas but the thresholded SDM data did not (Fig. 5). This may arise from a similar phenomenon to the above example, with the thresholded data removing information about large numbers of taxa with low but non-negligible probabilities of occurring there.

We also found signs of higher variance in the thresholded SDM results. This was particularly visible in low-diversity areas along the edges of the Central Valley and Los Angeles Basin, where the thresholded analysis classified individual isolated pixels as areas of significant endemism but the continuous data did not (Fig. 5). After thresholding, a very small number of taxa are left to drive the results in such sites, leading to high variance in observed diversity values and thus an increased chance of sites being “randomly” misidentified as areas of significant phylogenetic endemism. In contrast, the continuous SDM analysis includes data for many taxa in these low-diversity sites, which reduces the variance in the observed values, making “type 1 errors” less common. In short, because the continuous data contain more information, hypothesis tests on these data have reduced rates of false positives.

The second area of information loss that we addressed was in interpreting and presenting the results of statistical significance tests. Our results illustrate that by using continuous color ramps instead of discrete categories to represent p-values (Fig. 5), significance results can be communicated in a more informative and nuanced way. Just as it is best practice to report continuous p-values in written results, the same should be true for visual maps. Rather than simply classifying sites as significant or not based on an arbitrarily chosen alpha threshold, displaying maps of continuous p-values, particularly using a logit scale representing its order of magnitude, can communicate the degree of confidence in a given result. Using this approach, patterns within areas of endemism become more evident, highlighting the fact that arbitrary cut-offs tend to hide information that might be of biological relevance. While the choice to represent p-values as a gradient doesn’t involve a change in the methodology per se, it better represents the continuous nature of diversity patterns.

Finally, we also explored continuous alternatives to classifying sites into discrete biogeographic regions. Discrete categories may fit some data sets well, such as study systems in which a small number of relatively homogeneous regions are separated by distinct biogeographic breaks. But for the California bryophytes, as for many other systems, we found that distinct categories did not fit the data well. This reflects the fact that bryophyte diversity is distributed along relatively smooth gradients rather than falling into distinct zones. Using the combination of NMDS community ordination and a hierarchical dendrogram that was never “cut” into a fixed number of clusters, we were able to paint a richer and more accurate picture of bryophyte biogeography across the state (Fig. 3). While neither the ordination nor the dendrogram preserve the full dimensionality of the community turnover data, together they provide far more information than does a flat cluster analysis.

## Conclusion

Overall, mosses and liverworts share most regions of high endemism across California, with some interesting exceptions. Regions dominated by paleo-endemism are concentrated in coastal areas, which maintain more stable temperature and humidity across daily, annual, and millennial timescales due to the moderating influence of the Pacific Ocean. This is consistent with the biology of the majority of bryophytes, which are adapted to live in moist environments. On the other hand, regions dominated by neo-endemism are located in more interior areas, particularly east of the Sierra Nevada, which is a drier region due to the rain shadow effect. This observation is consistent with a relatively recent and perhaps ongoing diversification of some lineages in dry environments. Given the urgency of conservation needs and the limited availability of biodiversity data, particularly for understudied groups like bryophytes, studies should strive to make the most of available data. The threshold-free phylodiversity metrics we describe in this paper are a valuable set of tools to help achieve this. Additional approaches demonstrated in our analyses can also be useful, including judicious use of SDMs to augment sparse occurrence data, and the use of quantitative data on the degree of biodiversity protection offered by different land management zones across the state. Wider adoption of these methods could improve future studies focused on both basic biogeographic research and applied conservation and management.

## Acknowledgements

This work was possible thanks to the work of many individuals who collected, identified, recorded, sequenced, and made their specimen and DNA data publicly available. We are grateful for their efforts. MMK was supported by NSF grant #2019470. IGR was supported by a UC Mexus-CONACyT fellowship (number 709967).

## Author contributions

**Matthew M. Kling:** Conceptualization, Methodology, Formal Analysis, Writing – Original Draft, Visualization.

**Ixchel S. González-Ramírez:** Conceptualization, Formal Analysis, Data curation, Writing – Original Draft, Project Administration.

**Benjamin Carter:** Conceptualization, Data curation, Writing – Review and Editing, Supervision.

**Israel Borokini:** Conceptualization, Writing – Review and Editing.

**Brent Mishler:** Conceptualization, Writing – Review and Editing, Supervision.

## SUPPLEMENTARY MATERIALS

**Figure S1.**
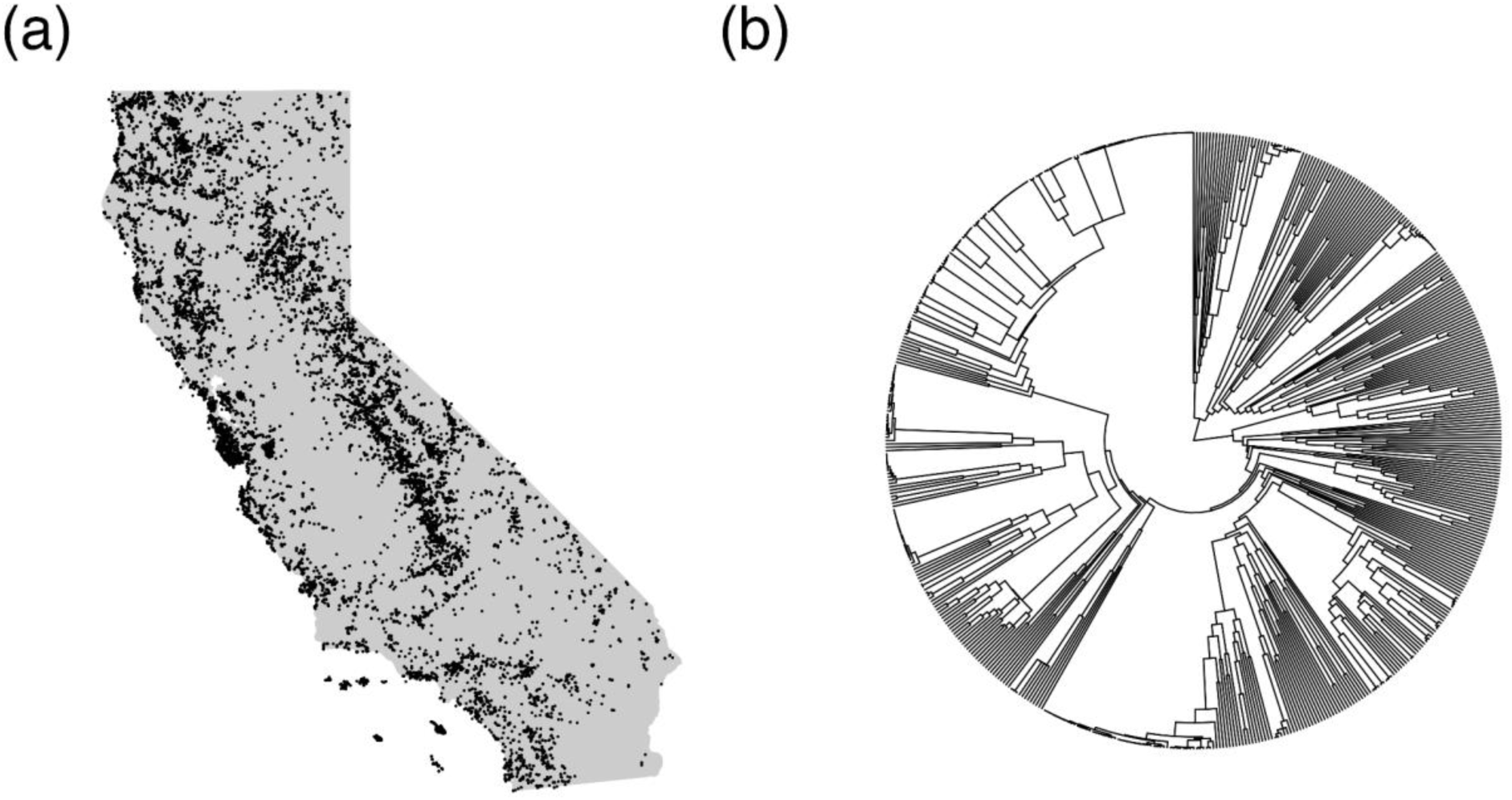
**(a)** Occurrence data and **(b)** phylogeny of California bryophytes used for this study.

**Figure S2.**
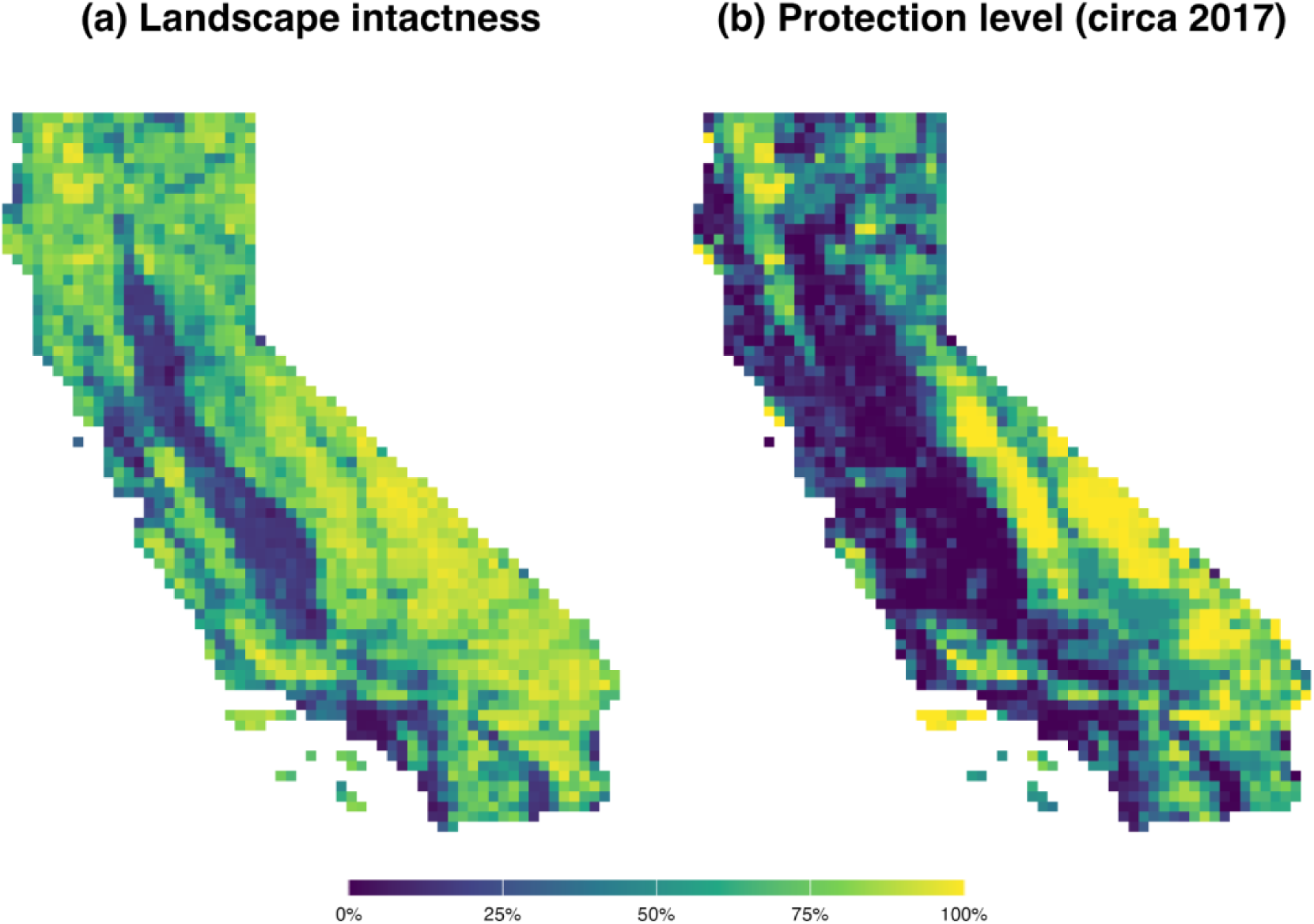
**(a)** Landscape intactness and **(b)** protection level of 15 km grid cells across California. Note that the analyses used landscape intactness at 1 km resolution.

**Figure S3.**
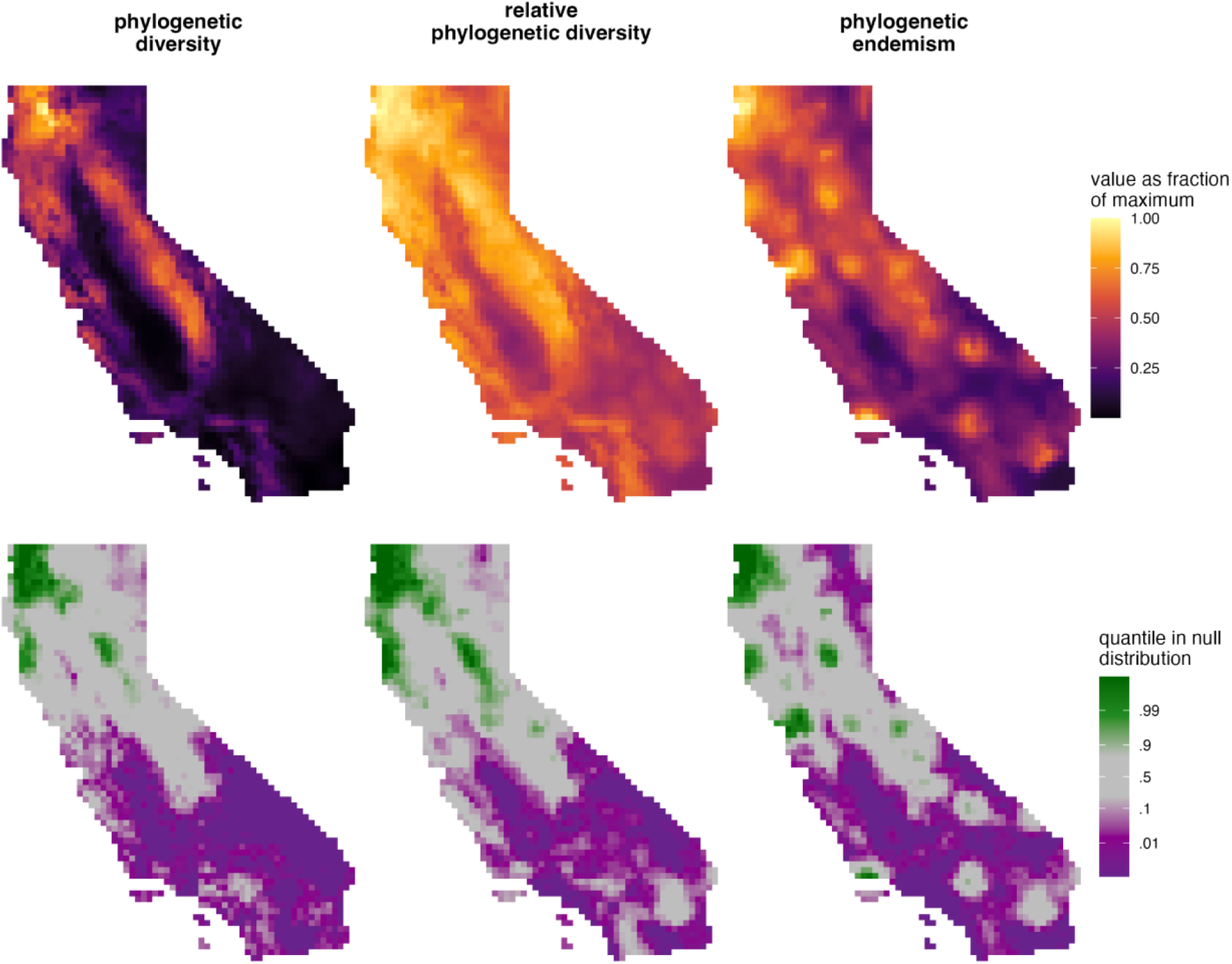
Alpha phylogenetic diversity measures for the mosses alone.

**Figure S4.**
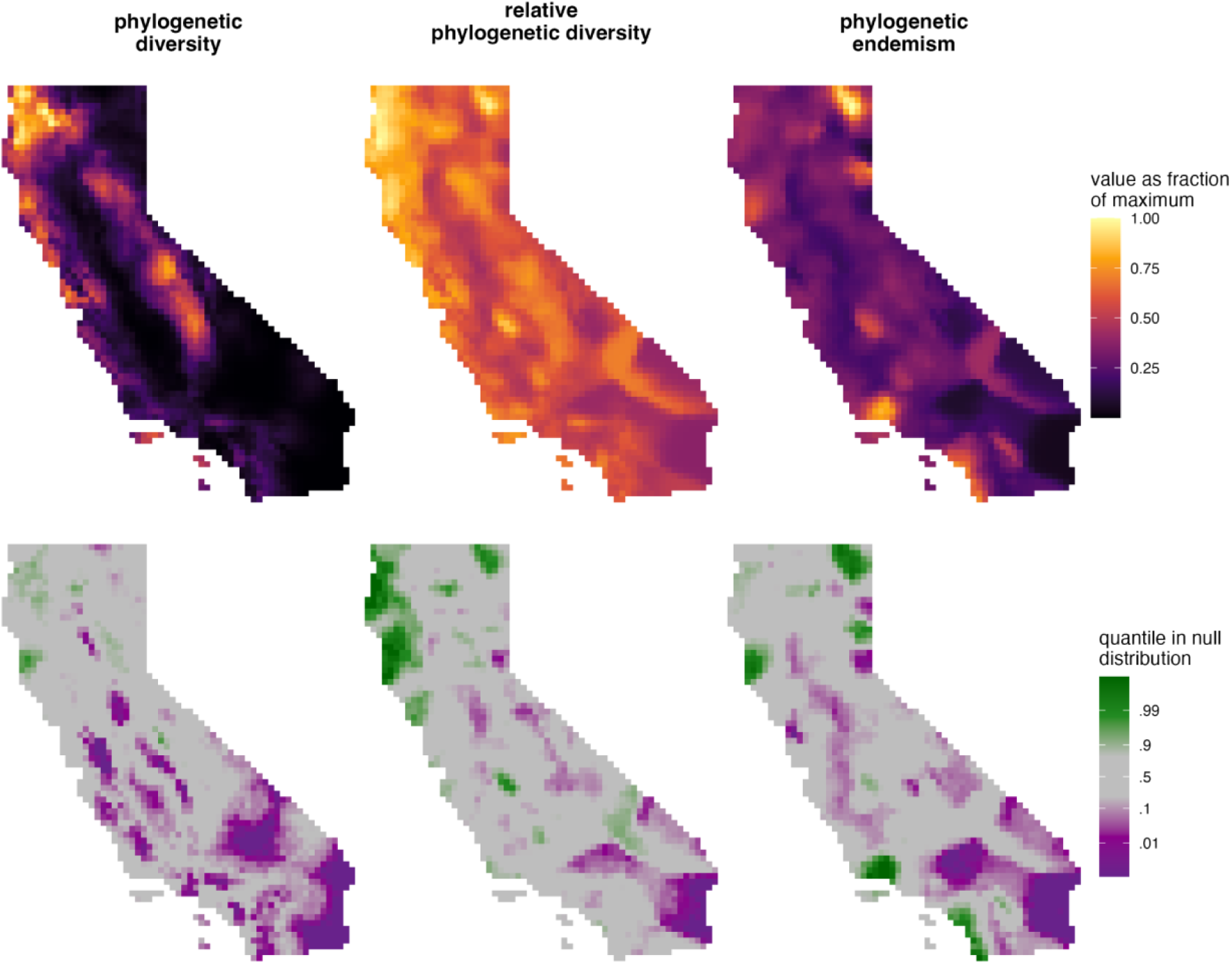
Alpha phylogenetic diversity measures for the liverworts alone.

**Figure S5.**
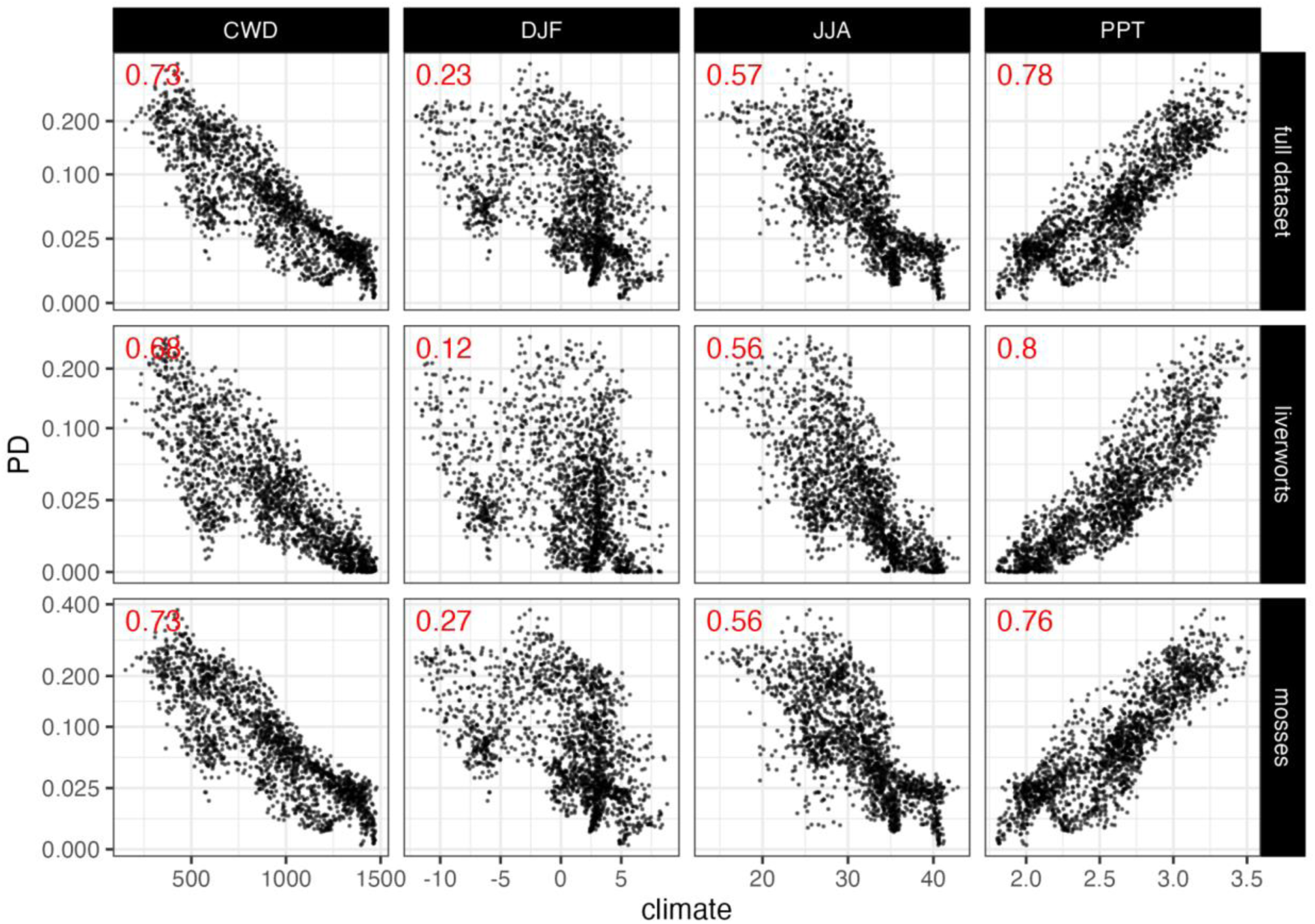
Relationships between macroclimate variables and PD for the full dataset, and for the mosses and liverworts alone. Relationships are shown for the four climate variables used to model species distributions: CWD (climatic water deficit, in mm), DJF (average winter nighttime minimum temperature, in degrees Celsius), JJA (average summer daytime maximum temperature, in degrees Celsius), and PPT (total annual precipitation, in log10 mm). Squared Spearman’s rank correlations for each pairwise relationship are shown in red.

**Table S1.** Calibration nodes selected from Betcheler et al., 2023. The uncalibrated rooted trees are available in the data repository.

| Node | Node number in uncalibrated rooted tree | Minimum age (My) | Maximum age (My) |
| --- | --- | --- | --- |
| Liverworts |  |  |  |
| Root | 120 | 444 | 450 |
| Complex thalloids | 121 | 387 | 395 |
| Sphaerocarpales | 123 | 272 | 287 |
| Aytoniaceae | 134 | 169 | 186 |
| Porellales | 167 | 330 | 349 |
| MRCA of *Marsupella* and *Blephanostora* | 219 | 213 | 225 |
| Mosses |  |  |  |
| Root | 467 | 416 | 423 |
| MRCA of *Psycomitrium* and *Discelium* | 508 | 245 | 258 |
| MRCA of *Orthodontium* and *Homalothecium* | 810 | 135 | 220 |
| MRCA of *Plagiomnium* and *Pohlia* | 712 | 58 | 116 |
| MRCA of *Tortula* and *Dicranowesia* | 617 | 155 | 213 |
| MRCA of *Grimmia* and *Blindia* | 540 | 163 | 193 |

**Table S2.**
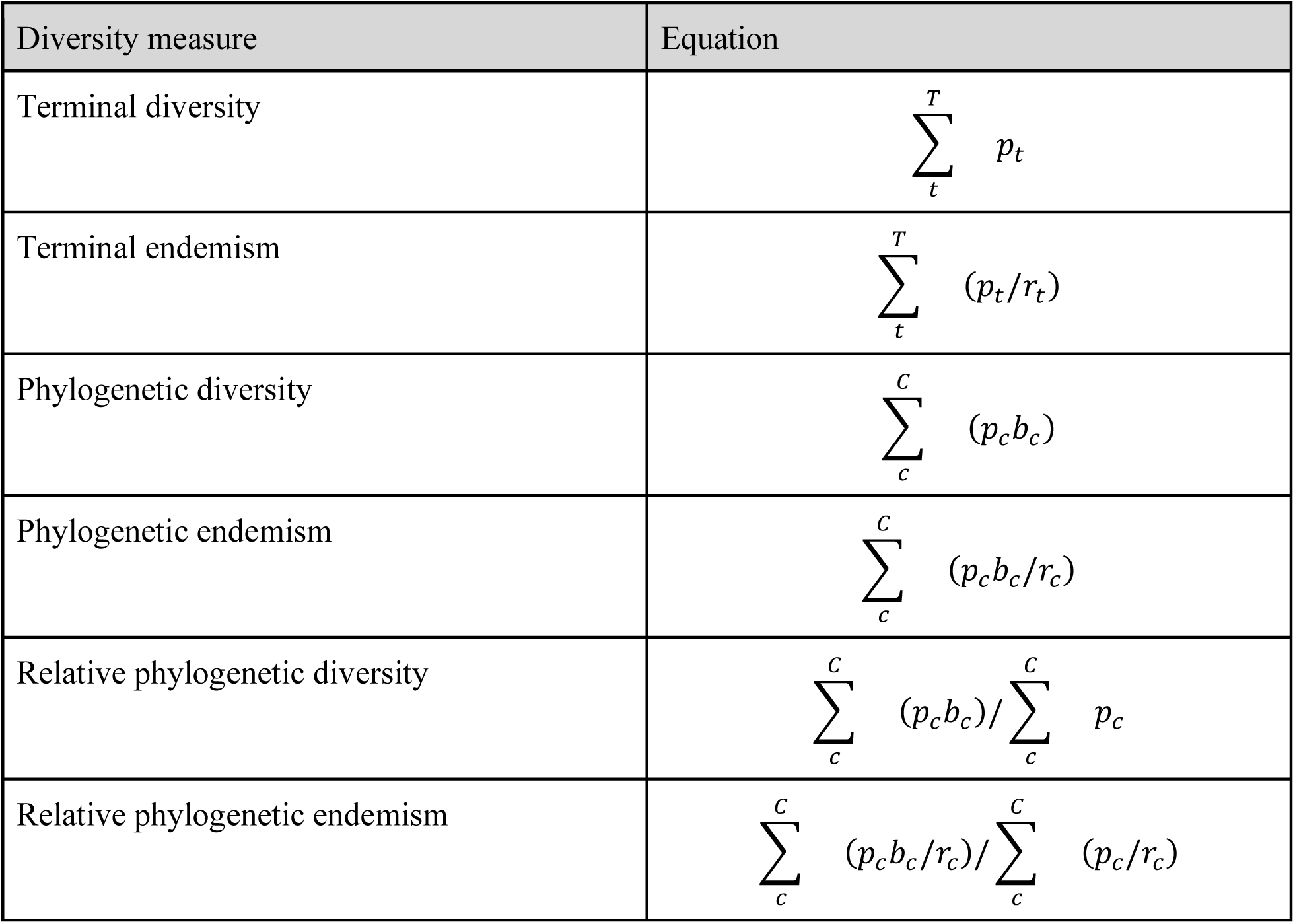
Equations for calculating alpha diversity measures using continuous probability data. In all equations, *t* indexes terminal taxa, *c* indexes clades (defined as nodes at all levels, including terminal and internal nodes), *p* is the probability of a clade occurring in a cell, *b* is the length of the subtending phylogenetic branch unique to a clade, and *r* is the clade’s range size (sum of *p* across all cells). *P_c_* is calculated as described in the main text.

